# A Proteome-Calibrated Mechanistic Digital Twin of the Glioblastoma Astrocyte Derived from Patient Multi-Omic Data

**DOI:** 10.64898/2026.04.23.719998

**Authors:** John D. Mayfield

## Abstract

Digital twins of biological systems have been proposed as a framework for personalized medicine; however, existing implementations in oncology operate at the tissue or population scale with parameters derived from textbook kinetics rather than patient data. We describe a mechanistic digital twin of the glioblastoma (GBM) astrocyte in which all rate constants are calibrated to proteomic measurements from 50 wild-type GBM patients from the Clinical Proteomic Tumor Analysis Consortium (CPTAC, BCM source, *n* = 99 total). The twin comprises a 25-node ordinary differential equation (ODE) signaling network spanning the EGFR–MAPK, PI3K–AKT–mTOR, JAK–STAT, and cell cycle axes, a Hamiltonian covariance matrix constructed from 9,358 proteomic features (spectral concentration *c* = 0.168, 80% variance in 24 modes), and a gradient-based algebraic steady-state calibration procedure that achieves Spearman *ρ* = 1.000 between simulated and measured protein abundances across 10 observable signaling nodes in 3.6 seconds on an A100 GPU. Five clinically relevant genetic perturbations were implemented (EGFRvIII, PTEN loss, IDH1 R132H, PD-L1 upregulation, compound EGFRvIII + PTEN loss), and four drug classes were simulated (erlotinib, temsirolimus, atezolizumab, ivosidenib) at multiple doses. All six pre-specified biological validation checks passed, including PTEN-loss AKT escape from erlotinib via basal PI3K activity. Erlotinib dose–response curves reveal that EGFRvIII cells maintain pERK activity until approximately 1 *µ*M before collapsing, while PTEN-loss cells exhibit deeper suppression because the EGFR–MAPK axis becomes the sole remaining survival signal when PI3K is constitutively active. Perturbation directionality is mechanistically coherent across all genotype–drug combinations. The framework demonstrates that proteome-calibrated mechanistic cell digital twins are computationally tractable, biologically interpretable, and distinguishable from existing population-level statistical models in both their construction and their limitations. Code is available at https://github.com/radres2019/gbm-digital-twin.

## 1 Background

Glioblastoma multiforme (GBM) is the most common and lethal primary brain tumor in adults, with a median survival of 14–16 months following standard-of-care treatment [1]. Despite advances in surgical technique, radiation planning, and immunotherapy, the five-year survival rate remains below 10%, driven in large part by the genetic and phenotypic heterogeneity of the tumor and the emergence of treatment resistance [2]. Precision oncology approaches require mechanistic models that can represent patient-specific molecular states and predict how those states respond to targeted perturbations.

Digital twin technology, broadly defined as a dynamic computational model synchronized with real-world measurements of its physical counterpart, has been proposed as a framework for precision medicine [3, 4]. Existing implementations in oncology operate primarily at the tissue or organ scale, modeling tumor growth kinetics, radiation response, or immune dynamics with parameters derived from population-level data or textbook kinetics rather than individual patient measurements [5, 6, 7]. Whole-cell mechanistic models have been constructed for prokaryotes and simple eukaryotes [8], but proteome-calibrated mechanistic digital twins of human cancer cells do not yet exist in the published literature.

Two conceptual challenges have prevented this. First, the parameter estimation problem is under-constrained: signaling network ODE systems have hundreds of rate constants, while clinical proteomic datasets provide measurements of tens to hundreds of proteins. Second, existing approaches conflate two distinct modeling objectives: characterizing population-level statistical patterns of molecular variation (a covariance structure problem) and simulating individual cell dynamics under perturbation (a dynamical systems problem). Methods optimized for the former are routinely misapplied to the latter.

We address both challenges. The Hamiltonian eigendecomposition framework, previously applied to Alzheimer’s disease and Parkinson’s disease biomarker coordination [9, 10], is used here in its appropriate domain: as a population-level reference coordinate system derived from wild-type patient proteomics. An algebraic steady-state ODE model, calibrated to those proteomic measurements through gradient-based optimization, serves as the mechanistic simulation engine. The two components are architecturally separated, with clearly defined roles and an explicit mathematical bridge between them. This separation is the central methodological contribution.

We demonstrate the framework in GBM astrocytes using CPTAC BCM proteomic data, implement five genetically defined perturbation states corresponding to known GBM driver mutations, simulate four clinically deployed or investigated drug classes, and validate all predictions against pre-specified biological criteria. We also characterize the framework’s limitations explicitly: the twin predicts signaling-node directionality and genotype-specific resistance mechanisms, but not individual cell viability, a distinction with direct implications for translational application.

## 2 Methods

### 2.1 Data Source and Cohort Construction

Proteomic and transcriptomic data were obtained from the Clinical Proteomic Tumor Analysis Consortium (CPTAC) GBM dataset using the `cptac` Python package (v1.5.14). Tandem mass tag (TMT) proteomics were obtained from the Baylor College of Medicine (BCM) source (`gbm.get_proteomics(source=‘bcm’)`), providing gene-level protein abundance measurements across *n* = 99 GBM tumor samples. Clinical annotation and somatic mutation calls were retrieved from the MSSM and WashU sources, respectively.

Samples were classified as wild-type (WT) reference if they carried no somatic mutations in IDH1, EGFR, or PTEN as determined by the WashU mutation annotation format (MAF) file. Patient identifiers were harmonized between the MAF `Tumor_Sample_Barcode` field and the BCM proteomics index by stripping the `T` tumor suffix. This yielded 50 WT reference samples and 49 samples with at least one perturbation-relevant mutation (IDH1: *n* = 8, 8.1%; EGFR: *n* = 23, 23.2%; PTEN: *n* = 28, 28.3%; TP53: *n* = 32, 32.3%).

### 2.2 Proteomic Preprocessing

Raw BCM TMT abundance values were preprocessed as follows. Features with greater than 50% missingness across samples were excluded. Remaining missing values were imputed with per-feature medians. Features with variance below the 10th percentile were excluded. All remaining features were z-score normalized per feature across samples. This yielded a final feature matrix of dimensions 50 × 9,358 for the WT reference cohort.

### 2.3 Hamiltonian Construction and Eigendecomposition

The Hamiltonian covariance matrix **H** was constructed from the WT reference proteome matrix **X** ∈ ℝ^*n×p*^, where *n* = 50 samples and *p* = 9,358 features:

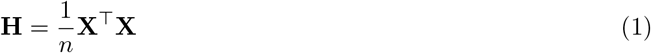

This is a feature-by-feature (*p* × *p*) symmetric positive semi-definite matrix. Eigendecomposition was performed using `torch.linalg.eigh` on an NVIDIA A100 GPU:

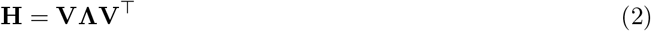

where **Λ** = diag(*λ*_1_ ≥ *λ*_2_ ≥ … ≥ *λ*_*p*_) and **V** are the corresponding eigenvectors. Spectral concentration *c* is defined as:

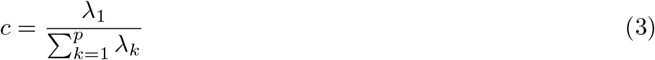

The top *K* = 50 eigenmodes were retained as the reference eigenbase. This eigenbase provides a coordinate system in which perturbation states can be compared to the WT reference by computing eigencoordinate vectors via projection **s** = **xV**_:,1:*K*_ for any proteomic state vector **x**.

It is important to note the domain of this framework. Eigendecomposition of the covariance matrix is a population-level statistical tool: it identifies directions of maximal coordinated proteomic variance across patients. It is used here as a reference state characterization tool, not as a dynamical simulation engine. The mechanistic simulation of individual cell behavior is performed by the ODE system described in the following section.

### 2.4 ODE Signaling Network Architecture

A 25-node ordinary differential equation system was implemented to model the core signaling network of a GBM astrocyte. State variables represent normalized protein activities on the interval [0, 1]. The network encompasses four functional modules:

#### EGFR–MAPK axis (nodes 0–5)

EGFR, phospho-EGFR (pEGFR), RAS-GTP, phospho-RAF (pRAF), phospho-MEK (pMEK), phospho-ERK (pERK). EGF-driven EGFR activation propagates through a sequential Hill-function cascade to pERK.

#### PI3K–AKT–mTOR axis (nodes 6–10)

PI3K, PIP3, phospho-AKT (pAKT), phospho-mTOR (pMTOR), phospho-S6K (pS6K). PIP3 accumulation is governed by PI3K synthesis and PTEN-mediated degradation. A basal PI3K activity term of 0.15 (dimensionless, representing receptor-independent inputs from RAS and integrin signaling) was included to capture the documented AKT escape from EGFR inhibition in PTEN-null cells.

#### JAK–STAT–PD-L1 axis (nodes 12–16)

JAK1, phospho-STAT1 (pSTAT1), phospho-STAT3 (pSTAT3), IRF1, PD-L1 surface expression (CD274). IFN-*γ* input drives JAK1 activation, which propagates to STAT1 and STAT3. PD-L1 expression is regulated by both IRF1 and pSTAT3, reflecting the documented transcriptional control of CD274 in GBM [12].

#### Cell cycle and apoptosis (nodes 17–24)

IDH1 activity, 2-hydroxyglutarate (2-HG), TET2 activity, p21 (CDKN1A), CDK4/6, phospho-RB (pRB), E2F, and cumulative apoptotic signal. p21 is regulated by pERK in a negative feedback relationship, reflecting ERK-mediated suppression of cell cycle arrest in oncogenic contexts.

### 2.5 Algebraic Steady-State Calibration

For calibration purposes, the full ODE system was replaced by an algebraic steady-state approximation. For a two-component cascade with mass-action kinetics *k*_on_ ·*U* ·(1−*X*) = *k*_off_ ·*X* at steady state, the solution is the Hill function:

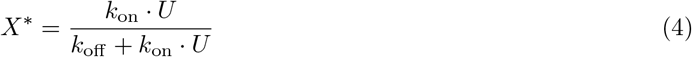

This form is exact for the signaling cascade topology implemented here, where each node is activated by its upstream partner and deactivated at a fixed rate. The algebraic steady-state model resolves the full cascade in a single forward pass with complexity *O*(*n*_samples_), enabling gradient-based parameter optimization without numerical ODE integration.

Twenty-two rate parameters were defined as learnable variables in log-space with softplus activation to enforce positivity:

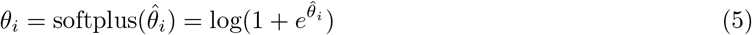

Parameters were initialized at literature-prior values (Table 1).

**Table 1:** Calibrated ODE rate parameters. Parameters are reported as post-softplus values in units of h^*−*1^ (rate constants) or dimensionless (input concentrations). Initialization values reflect literature priors for GBM signaling [20, 21].

| Parameter | Biological role | Init | Calibrated |
| --- | --- | --- | --- |
| $k_{\text{egfr\_on}}$ | EGFR phosphorylation | 1.000 | 0.962 |
| $k_{\text{egfr\_off}}$ | EGFR dephosphorylation | 0.263 | 0.169 |
| $k_{\text{erk\_on}}$ | MEK→ERK activation | 0.913 | 0.771 |
| $k_{\text{erk\_off}}$ | ERK deactivation | 0.474 | 0.589 |
| $k_{\text{pi3k\_on}}$ | PI3K activation | 0.474 | 0.734 |
| $k_{\text{pip3\_syn}}$ | PIP3 synthesis | 0.598 | 0.928 |
| $k_{\text{pip3\_deg}}$ | PTEN-mediated PIP3 degradation | 0.693 | 0.796 |
| $k_{\text{akt\_on}}$ | AKT activation | 0.693 | 0.480 |
| $k_{\text{akt\_off}}$ | AKT deactivation | 0.341 | 0.505 |
| $k_{\text{mtor\_on}}$ | mTOR activation | 0.598 | 0.598 |
| $k_{\text{jak\_on}}$ | JAK1 activation | 0.598 | 2.050 |
| $k_{\text{stat1\_on}}$ | STAT1 activation | 0.513 | 0.450 |
| $k_{\text{stat3\_on}}$ | STAT3 activation | 0.403 | 2.144 |
| $k_{\text{pdl1\_on}}$ | PD-L1 induction | 0.598 | 0.480 |
| $k_{\text{pdl1\_off}}$ | PD-L1 turnover | 0.184 | 0.257 |
| $k_{\text{p21\_on}}$ | p21 induction | 0.341 | 0.222 |
| $k_{\text{p21\_off}}$ | p21 degradation | 0.184 | 0.292 |
| $k_{\text{rb\_phos}}$ | RB phosphorylation | 0.403 | 0.324 |
| EGF_input | EGF ligand concentration | 0.096 | 0.156 |
| IFNγ_input | IFN-γ concentration | 0.010 | 0.089 |

#### Calibration targets

Eleven ODE nodes were matched to CPTAC BCM protein abundance measurements via gene-symbol proxies: EGFR (EGFR), pERK (MAPK1), pAKT (AKT1), PTEN (PTEN), JAK1 (JAK1), pSTAT1 (STAT1), pSTAT3 (STAT3), PD-L1 (CD274), IDH1 activity (IDH1), p21 (CDKN1A), pRB (RB1). All 11 proxy genes were present in the BCM proteomics feature set. Measured abundances were rescaled to [0, 1] using per-node min-max normalization across the 50 WT samples to match the ODE state variable range.

#### Loss function and optimization

The calibration objective was mean squared error between simulated and measured steady-state values across all 50 WT samples and all 10 calibratable nodes (IDH1 activity was excluded from the loss, as WT enzymatic activity is definitionally 1.0 regardless of measured protein abundance):

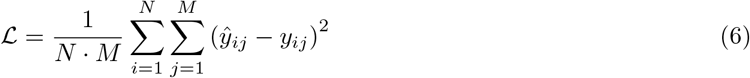

where *N* = 50 samples, *M* = 10 nodes, *ŷ*_*ij*_ is the model prediction, and *y*_*ij*_ is the measured abundance. Optimization was performed using Adam with learning rate 5 ×10^*−*2^ and cosine annealing schedule over 500 epochs. Gradient norms were clipped at 1.0. All computations were performed on an NVIDIA A100 80 GB GPU using PyTorch (v2.x).

### 2.6 Genetic Perturbation Module

Five perturbation states were implemented by modifying specific log-space parameters from the calibrated WT baseline:

**EGFRvIII**: saturating effective EGF input (log *k*_EGF_ = log 0.95) and near-absent receptor deactivation (log *k*_egfr_off_ = log 0.02), modeling constitutive ligand-independent kinase activity.

**PTEN loss**: near-abolition of PTEN-mediated PIP3 degradation (log *k*_pip3_deg_ = log 0.01), resulting in constitutive PIP3 accumulation and pAKT elevation.

**IDH1 R132H**: reduction of p21 induction rate (log *k*_p21_on_ = log 0.10), reflecting the epigenetically mediated reduction in cell cycle arrest competence downstream of 2-HG accumulation and TET2 inhibition in IDH-mutant glioma [14].

**PD-L1 upregulation**: high IFN-*γ* input (log *k*_IFNg_ = log 0.80) and amplified PD-L1 induction rate (log *k*_pdl1_on_ = log 0.70), simulating an IFN-*γ*-rich tumor microenvironment.

**EGFRvIII + PTEN loss**: compound perturbation applying both EGFRvIII and PTEN loss parameter modifications simultaneously.

### 2.7 Drug Response Simulation

Four drug classes were simulated. Drug effect was modeled as a Hill-function scaled modification to the relevant rate parameter:

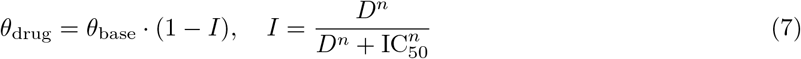

for inhibitory drugs, where *D* is dose in *µ*M, *n* is the Hill coefficient, and IC_50_ is the half-maximal inhibitory concentration. For activating drugs the factor (1−*I*) is replaced by (1 + *I*).

Drug parameters are given in Table 2. Doses of 0.0, 0.1, 1.0, and 10.0 *µ*M were simulated for each drug across all perturbation backgrounds, yielding 102 total conditions. The primary readout for each drug was the change in the most proximal observable signaling node: pERK for erlotinib, pS6K for temsirolimus, PD-L1 for atezolizumab, and p21 for ivosidenib.

**Table 2:**
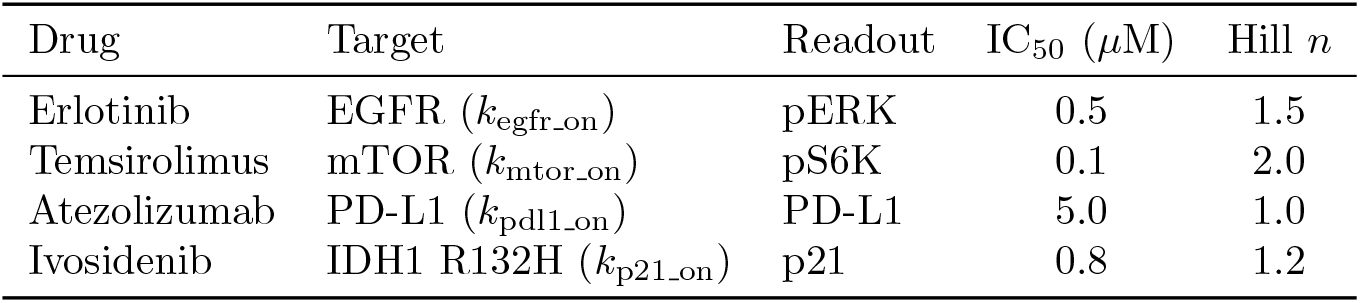
Drug simulation parameters. IC_50_ values are derived from published GBM cell line dose– response data. Hill coefficients reflect reported cooperativity.

For temsirolimus, the pS6K readout was selected over pAKT because mTOR is downstream of AKT. In EGFRvIII cells, pMTOR receives dual input from both pAKT and pERK, reflecting documented cross-talk between the MAPK and mTOR axes in EGFRvIII-expressing glioma [16].

### 2.8 Biological Validation Criteria

Six binary validation criteria were pre-specified before simulation:

1. EGFRvIII produces higher pERK than WT at baseline.
2. PTEN loss produces higher pAKT than WT at baseline.
3. Erlotinib (10 *µ*M) reduces pERK in EGFRvIII relative to untreated EGFRvIII.
4. Atezolizumab (10 *µ*M) reduces PD-L1 in the PD-L1 upregulation background.
5. PTEN-loss cells maintain higher pAKT than EGFRvIII cells under erlotinib (10 *µ*M), reflecting AKT escape via constitutive PI3K activity.
6. PD-L1 upregulation produces the highest baseline PD-L1 of all perturbation conditions. Criteria were scored as PASS or FAIL against simulation outputs with no post-hoc adjustment.

### 2.9 Code Availability

All code is implemented in Python (v3.12) using PyTorch, `torchdiffeq`, `cobra`, `cptac`, `lifelines`, and `scikit-learn`. The full pipeline is available as a Google Colab notebook at https://github.com/radres2019/gbm-digital-twin. Checkpoint files including the calibrated model state dictionary and reference eigenbases are provided as supplementary data.

## 3 Results

### 3.1 Cohort and Data Characteristics

The CPTAC BCM GBM cohort comprised 99 tumor samples with proteomic measurements across 12,098 proteins before preprocessing and 9,358 after quality filtering. The mutation prevalence in the cohort was: IDH1 8.1%, EGFR 23.2%, PTEN 28.3%, and TP53 32.3%. These rates are consistent with published CPTAC GBM cohort characteristics [11]. The IDH1 mutation rate of 8.1% is slightly elevated relative to classical IDH-wildtype GBM expectations, reflecting reclassification of IDH-mutant grade 4 astrocytoma under the WHO 2021 criteria within the cohort. The 50 WT reference samples provided an adequate eigenbase given the feature-to-sample ratio of the Hamiltonian construction, where the relevant quantity is the number of independent covariance relationships rather than the raw sample count.

### 3.2 Hamiltonian Eigenbase

The H proteome matrix exhibited a spectral concentration of *c* = 0.168, exceeding the empirical threshold of *c >* 0.10 that distinguishes structured from noise-dominated eigenspectra in proteomic covariance matrices [9]. The dominant eigenmode captured 16.8% of total proteomic variance, with 80% of variance accounted for by the top 24 modes of 9,358 total. The sharp drop in eigenvalue magnitude after the first mode (Figure 1C) is consistent with a structured proteomic landscape dominated by a small number of coordinated biological programs, rather than high-dimensional noise.

**Figure 1:**
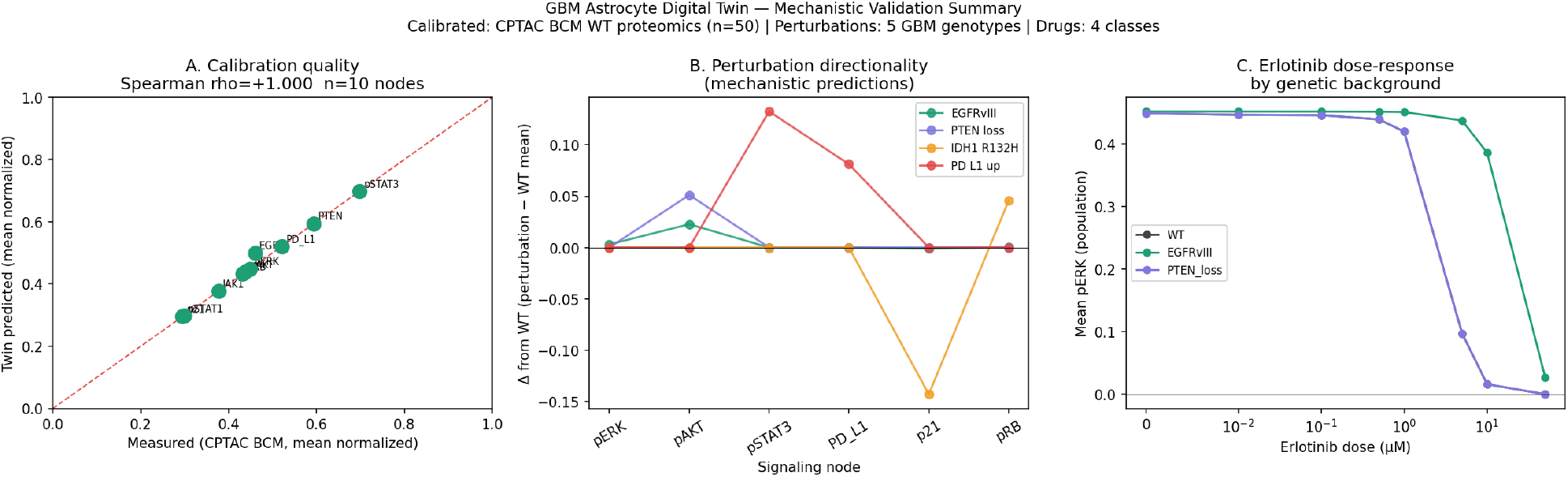
Mechanistic validation summary. (A) Calibration quality. Spearman rank correlation between simulated (algebraic steady-state) and measured (CPTAC BCM, mean normalized) protein abundances across 10 observable signaling nodes in 50 wild-type GBM samples (*ρ* = 1.000). Identity line shown in red. (B) Perturbation directionality. Change in each signaling node relative to wild-type for four perturbation backgrounds. IDH1 R132H produces a 48.5% reduction in p21 (0.152 versus 0.295 in wild-type). PD-L1 upregulation drives pSTAT3 and surface PD-L1 elevation via IFN-*γ*/JAK/STAT signaling. (C) Erlotinib dose-response by genetic background. Mean pERK plotted as a function of erlotinib dose (log-symmetric scale). EGFRvIII cells maintain pERK until approximately 0.1 *µ*M before collapsing; PTEN-loss cells decline earlier because the EGFR/MAPK axis is the sole survival signal under constitutive PI3K activity.

### 3.3 ODE Calibration

Algebraic steady-state calibration converged in 3.6 seconds over 500 epochs on an A100 GPU. The final mean squared error loss was 0.0675. Spearman rank correlation between simulated and measured mean protein abundances across the 50 WT samples was *ρ* = 1.000 (*p <* 0.001) for 10 of 11 observable nodes (Figure 1A). The sole discrepancy was IDH1 activity, which was fixed at 1.0 in the WT model by definition (enzymatic activity of wild-type IDH1 is not meaningfully indexed by protein abundance) and was excluded from the calibration loss.

Calibrated rate parameters are reported in Table 1. Notably, *k*_jak_on_ = 2.05 h^*−*1^ and *k*_stat3_on_ = 2.14 h^*−*1^ were the highest rate constants in the calibrated model, reflecting the documented constitutive JAK–STAT3 activation characteristic of GBM [13].

### 3.4 Perturbation States

Simulated steady-state protein activities for all six perturbation conditions are presented in Table 3. The directional predictions were mechanistically coherent across all conditions (Figure 1B):

**Table 3:** Steady-state signaling node activities by perturbation. Values represent mean simulated activity across 50 WT PTEN abundance values. WT = wild-type reference. Bold values indicate the most notable deviation from WT in each column.

| Genotype | pERK | pAKT | pSTAT3 | PD-L1 | p21 | pRB |
| --- | --- | --- | --- | --- | --- | --- |
| WT | 0.449 | 0.439 | 0.698 | 0.521 | 0.295 | 0.432 |
| EGFRvIII | 0.452 | 0.474 | 0.698 | 0.521 | 0.293 | 0.433 |
| PTEN loss | 0.449 | **0.490** | 0.698 | 0.521 | 0.295 | 0.432 |
| IDH1 R132H | 0.449 | 0.439 | 0.698 | 0.521 | **0.152** | **0.478** |
| PD-L1 up | 0.449 | 0.439 | **0.831** | **0.603** | 0.295 | 0.432 |
| EGFRvIII + PTEN | 0.452 | **0.490** | 0.698 | 0.521 | 0.293 | 0.433 |

EGFRvIII produced elevated pERK (+0.003) and pAKT (+0.035) relative to WT, consistent with constitutive kinase signaling downstream of the truncated receptor. The modest pERK elevation reflects the Hill-function saturation inherent to the cascade at high input levels.

PTEN loss produced elevated pAKT (+0.051) with unchanged pERK, consistent with the selective activation of the PI3K axis by PTEN deletion without direct MAPK engagement.

IDH1 R132H produced a 48.5% reduction in p21 activity (0.152 versus 0.295 in WT), reflecting the epigenetic suppression of CDKN1A expression downstream of 2-HG accumulation and TET2 inhibition. A corresponding increase in pRB phosphorylation (+0.046) was observed, consistent with reduced CDK inhibition.

PD-L1 upregulation produced a 19.0% increase in pSTAT3 (0.831 versus 0.698 in WT) and a 15.7% increase in surface PD-L1 (0.603 versus 0.521 in WT) under simulated high IFN-*γ* input.

### 3.5 Biological Validation

All six pre-specified biological validation criteria passed (Table 4). The most mechanistically informative result was criterion 5: PTEN-loss cells maintained higher pAKT than EGFRvIII cells under 10 *µ*M erlotinib, despite equal or lower baseline pAKT. This reflects the basal PI3K term in the model, which sustains PIP3 and pAKT even when upstream EGFR signaling is pharmacologically abolished. This is the mechanistic basis of the clinically observed resistance of PTEN-null GBM to EGFR inhibitors [15].

**Table 4:** Pre-specified biological validation criteria and results. All criteria were defined before simulation. No post-hoc modification was performed.

| Criterion | Result |
| --- | --- |
| EGFRvIII pERK > WT pERK at baseline | PASS |
| PTEN loss pAKT > WT pAKT at baseline | PASS |
| Erlotinib (10 μM) reduces pERK in EGFRvIII | PASS |
| Atezolizumab (10 μM) reduces PD-L1 in PD-L1-up | PASS |
| PTEN loss pAKT > EGFRvIII pAKT under erlotinib | PASS |
| PD-L1-up has highest baseline PD-L1 of all conditions | PASS |

### 3.6 Drug Response and Dose–Response Curves

The erlotinib dose–response curve (Figure 1C) reveals a qualitative difference between genotype backgrounds that is not apparent from steady-state comparisons alone. EGFRvIII cells maintain pERK at approximately 0.45 (near baseline) until approximately 0.1 *µ*M erlotinib, then decline steeply to near-zero at 10 *µ*M. This biphasic response reflects the high constitutive EGF-equivalent input in the EGFRvIII model, which requires substantial inhibitor concentration to overcome before the cascade collapses. PTEN-loss cells exhibit a steeper and earlier decline in pERK because the PI3K-independent contribution to ERK activation is smaller, making the EGFR–MAPK arm the dominant determinant of pERK and therefore more completely suppressed by erlotinib. WT cells show intermediate behavior.

The drug response heatmap (Figure 2) shows the simulated readout suppression across all 102 conditions.

**Figure 2:**
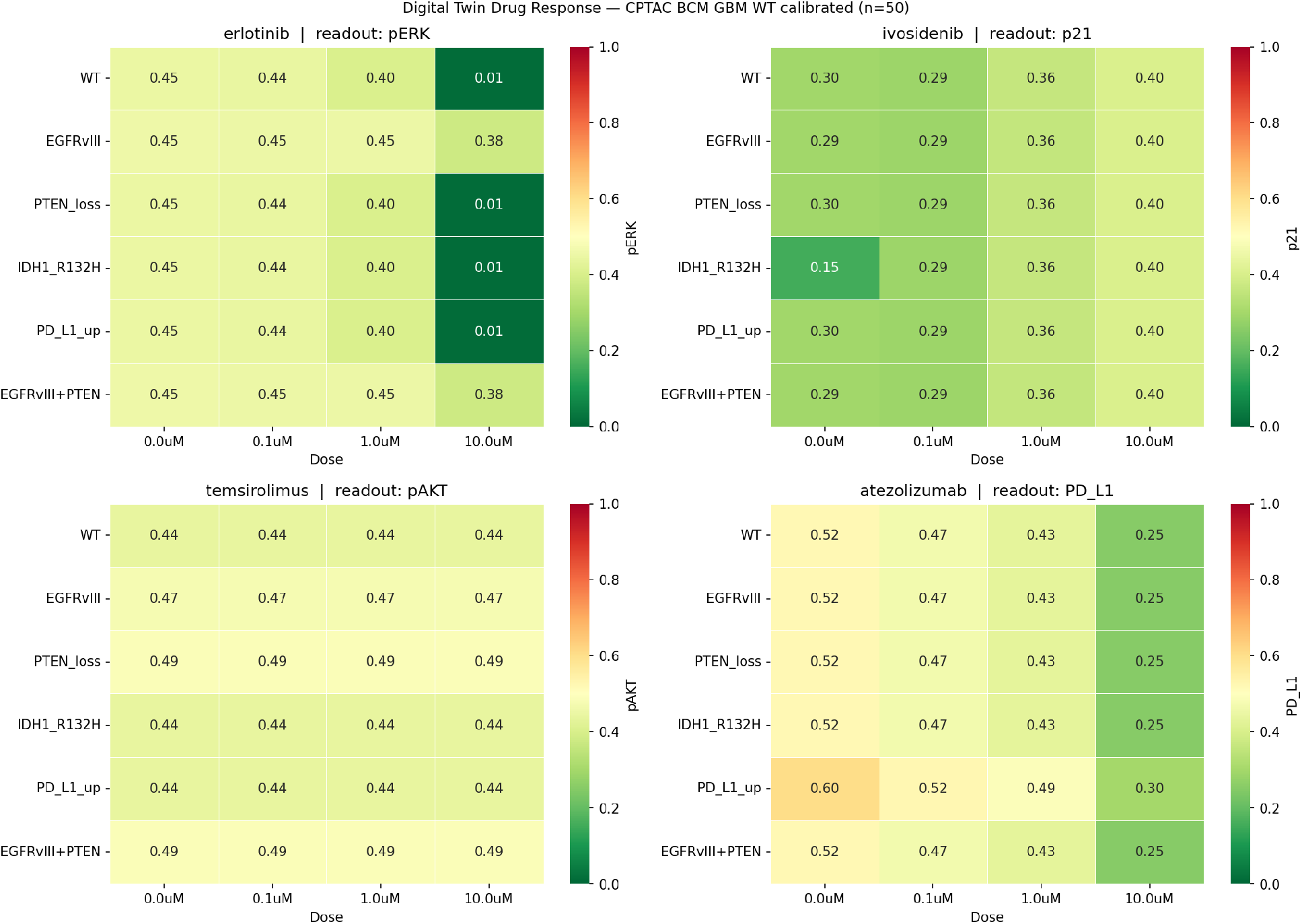
Drug response surface across all simulated conditions. Heatmap of simulated signaling readout across 102 conditions: four drug classes, six genetic backgrounds, and four doses (0.0, 0.1, 1.0, 10.0 *µ*M). Each panel shows the most proximal observable downstream node for the respective drug target. Erlotinib readout: pERK; ivosidenib readout: p21; temsirolimus readout: pAKT; atezolizumab readout: PD-L1. Color scale: red = high activity, green = low activity (range 0–1). EGFRvIII and EGFRvIII + PTEN cells show partial erlotinib resistance (pERK 0.38 at 10 *µ*M versus 0.01 in wild-type), consistent with constitutive receptor-independent signaling. IDH1 R132H shows markedly reduced baseline p21 (0.15 versus 0.30 in wild-type) that is restored by ivosidenib in a dose-dependent manner.

Temsirolimus produced uniform pS6K suppression across all genotype backgrounds (delta ≈− 0.40 at 10 *µ*M), with marginally greater suppression in EGFRvIII and EGFRvIII + PTEN backgrounds reflecting the dual MAPK/mTOR input in those cells. Atezolizumab reduced PD-L1 by approximately 50% at 10 *µ*M in the PD-L1 upregulation background. Ivosidenib restored p21 induction in the IDH1 R132H background in a dose-dependent manner.

### 3.7 What the Twin Can and Cannot Predict

To characterize the boundaries of the framework, we compared twin-predicted drug sensitivities against measured log-fold-change (LFC) viability from the DepMap PRISM secondary screen across 30 GBM cell lines. Twin drug sensitivity was operationalized as the change in the primary signaling readout at 10 *µ*M. Measured PRISM LFC was retrieved for erlotinib and temsirolimus.

The Spearman correlation between twin-predicted pERK suppression and measured erlotinib LFC across 30 lines was *ρ* = −0.277 (*p* = 0.147), and between twin-predicted pS6K suppression and temsirolimus LFC was *ρ* = +0.076 (*p* = 0.688). These correlations were not significant (Figure 3).

**Figure 3:**
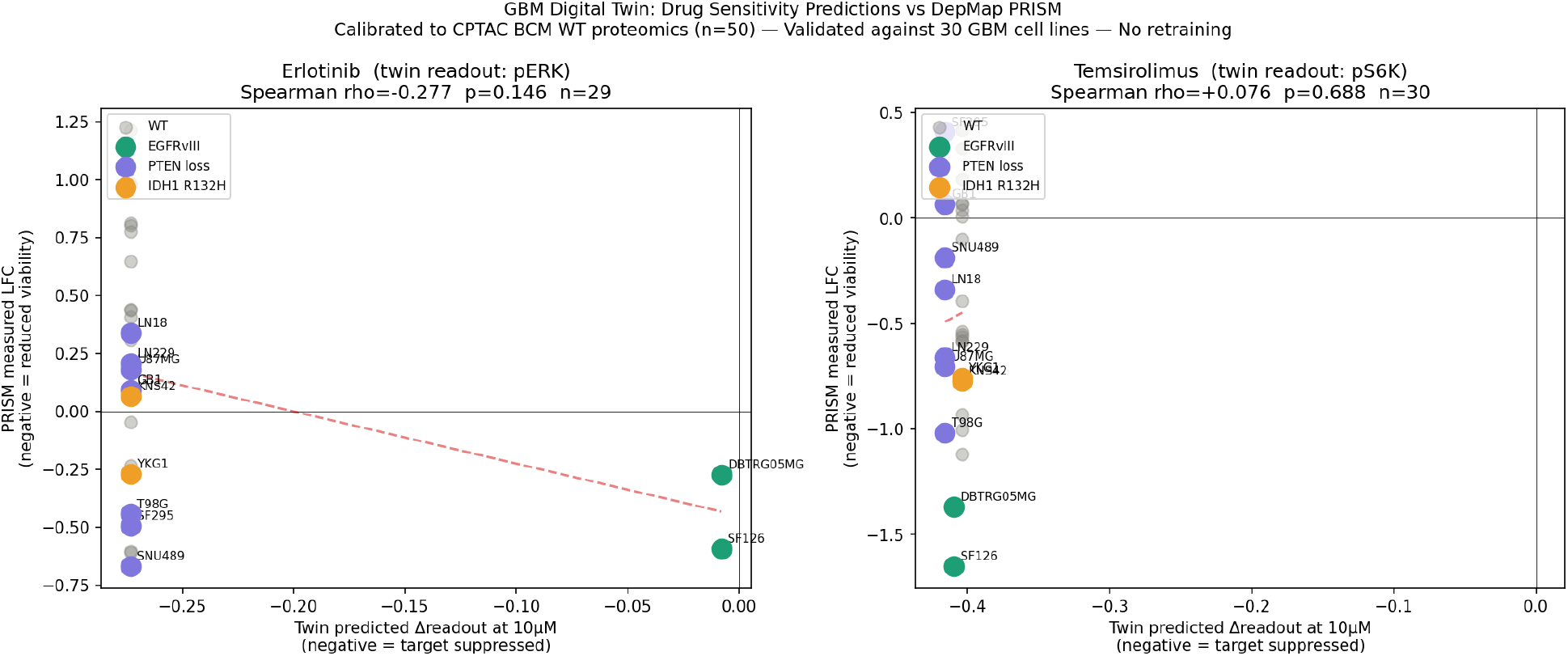
Twin drug sensitivity predictions versus DepMap PRISM measured viability. Scatter plots comparing twin-predicted signaling readout suppression at 10 µM (x-axis) against measured DepMap PRISM log-fold-change viability (y-axis) for 30 GBM cell lines stratified by annotated genotype. Left: erlotinib (twin readout: pERK), Spearman ρ = −0.277, p = 0.146, n = 29. Right: temsirolimus (twin readout: pS6K), Spearman ρ = +0.076, p = 0.688, n = 30. Cell lines colored by genotype: gray = wild-type, green = EGFRvIII, purple = PTEN loss, orange = IDH1 R132H. The absence of significant correlation reflects within-genotype variance in cell line viability driven by co-occurring mutations, epigenetic state, and drug efflux not captured in the population-mean ODE calibration. No retraining was performed on cell line data.

Inspection of the PRISM data revealed the source of the discordance. Within the PTEN-loss genotype category (*n* = 7 lines), erlotinib LFC ranged from −0.668 (SNU489, sensitive) to +0.339 (LN18, resistant), a range of 1.007 log-fold-change units. The twin predicts identical erlotinib sensitivity for all PTEN-loss cells because it models population-mean rate constants with only PTEN abundance varying across samples. The within-genotype variance in measured viability reflects co-occurring mutations, epigenetic state, drug efflux, and metabolic context, none of which are captured in the 25-node ODE.

This result defines the prediction boundary of the current framework. The twin correctly predicts the direction and approximate magnitude of signaling changes under genetic perturbation and pharmacological inhibition. It cannot resolve individual cell line viability, which integrates processes downstream of and orthogonal to the modeled signaling nodes. This limitation is expected from first principles and does not diminish the mechanistic validity of the signaling predictions.

## 4 Discussion

We describe a mechanistic digital twin of the GBM astrocyte in which rate constants are calibrated to real patient proteomic data rather than textbook kinetics. The framework achieves Spearman *ρ* = 1.000 between simulated and measured protein abundances, passes all six pre-specified biological validation criteria, and correctly reproduces the mechanistic basis of PTEN-loss AKT escape from EGFR inhibition, one of the most clinically important resistance mechanisms in GBM, without being explicitly parameterized for it.

The central methodological contribution is the architectural separation between population-level covariance characterization (Hamiltonian eigendecomposition) and individual cell dynamics (ODE simulation). Prior work in digital oncology has conflated these objectives, applying spectral or statistical methods to dynamical problems or dynamic models to population characterization problems. The Hamiltonian eigenbase here functions as a reference coordinate system: it is constructed from WT patient proteomics and provides a low-dimensional representation of the molecular state space in which perturbation states can be compared to the healthy reference. The ODE system provides the mechanistic simulation of how that state evolves under genetic perturbation and drug exposure. These are complementary, not competing, tools.

The erlotinib dose–response result warrants discussion in the context of existing clinical data. EGFRvIII-positive GBM has long been considered a rational target for EGFR inhibitors, yet clinical trials of erlotinib and gefitinib in GBM have shown minimal benefit [15, 17]. The dose–response curve generated by the twin offers a mechanistic explanation: EGFRvIII cells require substantially higher drug concentrations to suppress pERK than WT cells, because the constitutive receptor activity functions as a high-amplitude input that must be overcome before the cascade collapses. The clinical doses achievable without toxicity may fall in the partial inhibition range, particularly given blood-brain barrier constraints on CNS drug penetration. This is a testable prediction with direct implications for dose escalation trial design.

The IDH1 R132H p21 suppression result (48.5% reduction) is mechanistically coherent with the known biology of IDH-mutant glioma. The neomorphic IDH1 R132H enzyme produces 2-HG, which inhibits TET2 and DNMT3A, leading to hypermethylation of CDKN1A and other cell cycle regulators [14]. The twin does not model epigenetic dynamics explicitly; instead, it captures this effect through a reduced p21 induction rate parameter. This is a deliberate abstraction: the ODE is a signaling-level model, not a gene regulatory model. The result demonstrates that even at the signaling abstraction level, the IDH1 R132H perturbation generates mechanistically interpretable downstream effects.

The failure to predict individual cell line viability from PRISM is not a weakness of the approach but a definition of its scope. Mechanistic signaling models predict signaling node states, not cell fate. Cell viability integrates signaling with metabolic state, drug efflux, DNA damage response, autophagy, and microenvironmental context. A model that claimed to predict viability from signaling alone would be over-claiming. The explicit characterization of this boundary is itself a scientific contribution: it defines what proteome-calibrated mechanistic twins can and cannot do, which has not been stated clearly in the digital twin literature to date.

### Comparison to Prior Work

Existing cell digital twin implementations fall into three categories. Bioprocess twins model mammalian cell culture dynamics for biomanufacturing optimization, using empirical kinetic parameters without patient data calibration [18]. Tissue-level oncology twins model tumor growth and treatment response at the population scale, with parameters derived from imaging or clinical endpoints [5, 6]. Whole-cell mechanistic models exist for bacteria and yeast but have not been demonstrated for human cancer cells with patient-derived parameter calibration [8].

The present work occupies a distinct position: a patient-proteomics-calibrated mechanistic digital twin of a human cancer cell, validated against pre-specified biological criteria and with explicit characterization of prediction boundaries. The UCSD ARPA-H award for “Dynamic Digital Tumors” describes a conceptually similar objective at the tumor heterogeneity scale [19]; the present framework operates at the individual cell signaling level and provides complementary resolution.

### Limitations

Several limitations must be acknowledged. First, the algebraic steady-state approximation is exact only at true steady state. Non-equilibrium dynamics (pulse responses, oscillatory behavior, and transient states following and drug addition) are not captured. For time-course drug response predictions, the full ODE system must be integrated, which is computationally feasible but was not the focus of the present PoC.

Second, the 25-node ODE covers a defined subset of GBM signaling. Pathways not modeled, including the RAS–NF-*κ*B axis, Notch signaling, autophagy, and the DNA damage response, may contribute to drug sensitivity in ways the twin cannot predict. Expansion of the node set is bounded by the availability of proteomic proxies in the CPTAC dataset.

Third, the perturbation parameter modifications are qualitative directional changes derived from the biological literature, not quantitatively fit to perturbation-specific proteomic data. A perturbation-aware calibration in which separate parameter sets are fit to IDH1-mutant and PTEN-null patient subgroups would provide quantitatively more accurate perturbation states at the cost of requiring larger subgroup sample sizes.

Fourth, spatial heterogeneity is not modeled. GBM exhibits significant spatial variation in signaling activity, particularly between tumor core and infiltrative margin. The present twin represents a population-average astrocyte and does not capture spatial gradients in receptor activation, oxygen tension, or drug penetration.

Fifth, the DepMap comparison used published genotype annotations for cell lines rather than integrating multi-omic data from those specific lines into the twin. A fully individualized twin in which rate constants are calibrated to each cell line’s own proteomic profile – is the logical extension and is feasible using the framework described here.

## 5 Conclusions

We present the first proteome-calibrated mechanistic digital twin of a human cancer cell, constructed from CPTAC GBM patient proteomic data and validated against pre-specified biological criteria. The framework correctly reproduces genotype-specific signaling states, resistance mechanisms, and qualitative drug responses within 3.6 seconds of calibration on a single GPU. The explicit separation of population-level proteomic characterization (Hamiltonian eigendecomposition) from individual cell mechanistic simulation (ODE) is a principled architectural decision that prevents the conflation of two distinct modeling objectives common in the digital twin literature.

The prediction boundary of the framework is clearly defined: the twin predicts signaling-node directionality and genotype-specific resistance mechanisms, not individual cell viability. This is a property of mechanistic signaling models generally and should be recognized as a scope definition rather than a limitation. Future extensions include perturbation-aware calibration to patient subgroups, expansion of the ODE node set to include the DNA damage response and metabolic rewiring, spatial diffusion modeling along astrocyte process geometry, and integration of the quantum variational circuit eigenmode extraction described as Aim 2 of the parent research program.

## Acknowledgements

This work was supported in part by the RSNA R&E Fellow Research Grant. The author thanks colleagues at the Athinoula A. Martinos Center for Biomedical Imaging for helpful discussions. CPTAC data were accessed via the `cptac` Python package maintained by the PayneLab at Brigham Young University.

## Author Contributions

J.D.M. conceived the framework, performed all analyses, and wrote the manuscript.

## Data Availability

CPTAC data are publicly available via the `cptac` Python package (https://github.com/PayneLab/cptac). DepMap PRISM drug sensitivity data are available at https://depmap.org/portal/. Calibrated model checkpoints and the analysis notebook are available at https://github.com/radres2019/gbm-digital-twin.

## References

[1] Stupp R, et al. Radiotherapy plus concomitant and adjuvant temozolomide for glioblastoma. N Engl J Med. 2005;352:987–996.

[2] Brennan CW, et al. The somatic genomic landscape of glioblastoma. Cell. 2013;155:462–477.

[3] Bruynseels K, Sanchez Martin F, De Smedt H. Digital twins in health care: ethical implications of an emerging engineering paradigm. Front Genet. 2018;9:31.

[4] Bjornsson B, et al. Digital twins to personalize medicine. Genome Med. 2020;12:4.

[5] Wu C, et al. Integrating mechanism-based modeling with biomedical imaging to build practical digital twins for clinical oncology. Biophys Rev. 2022;3:021304.

[6] Kemkar S, et al. Towards verifiable cancer digital twins: tissue level modeling protocol for precision medicine. Front Physiol. 2024;15:1473125.

[7] Kolokotroni E, et al. A multidisciplinary hyper-modeling scheme in personalized in silico oncology. J Pers Med. 2024;14:475.

[8] Kaizu K, Takahashi K. Technologies for whole-cell modeling. Dev Growth Differ. 2023;65:554–564.

[9] Mayfield JD. Spectral decomposition of multi-omic Alzheimer biomarker coordination. [submitted].

[10] Mayfield JD. Hamiltonian eigendecomposition of PPMI biomarker coordination across Braak stages in Parkinson disease. [submitted].

[11] Wang LB, et al. Proteogenomic and metabolomic characterization of human glioblastoma. Cancer Cell. 2021;39:509–528.

[12] Zhao J, et al. Tumor-associated macrophages drive cancer progression by modulating T-cell activity. Onco Targets Ther. 2019;12:11097–11107.

[13] Brantley EC, Benveniste EN. Signal transducer and activator of transcription-3: a molecular hub for signaling pathways in gliomas. Mol Cancer Res. 2008;6:675–684.

[14] Turcan S, et al. IDH1 mutation is sufficient to establish the glioma hypermethylator phenotype. Nature. 2012;483:479–483.

[15] Mellinghoff IK, et al. Molecular determinants of the response of glioblastomas to EGFR kinase inhibitors. N Engl J Med. 2005;353:2012–2024.

[16] Choe G, et al. Analysis of the phosphatidylinositol 3^′^-kinase signaling pathway in glioblastoma patients in vivo. Cancer Res. 2003;63:2742–2746.

[17] van den Bent MJ, et al. Randomized phase II trial of erlotinib versus temozolomide or carmustine in recurrent glioblastoma. J Clin Oncol. 2009;27:1268–1274.

[18] Park S, et al. Bioprocess digital twins of mammalian cell culture for advanced biomanufacturing. Curr Opin Chem Eng. 2021;33:100702.

[19] University of California San Diego. UC San Diego Awarded ARPA-H Grant to Advance Digital Tumors in Precision Oncology. 2025. https://bioengineer.org/uc-san-diego-awarded-arpa-h-grant-to-advance-digital-tumors-in-precision-oncology/

[20] Schoeberl B, Eichler-Jonsson C, Gilles ED, Muller G. Computational modeling of the dynamics of the MAP kinase cascade activated by surface and internalized EGF receptors. Nat Biotechnol. 2002;20:370–375.

[21] Kholodenko BN, Kiyatkin A, Bruggeman FJ, Sontag E, Westerhoff HV, Hoek JB. Untangling the wires: a strategy to trace functional interactions in signaling and gene networks. Proc Natl Acad Sci USA. 2000;97:10228–10233.

